# Higher order interactions and coexistence theory

**DOI:** 10.1101/748517

**Authors:** Pragya Singh, Gaurav Baruah

**Affiliations:** Zoological Institute, Evolutionary Biology, University of Basel; Department of Evolutionary Biology and Environmental studies, University of Zurich

**Keywords:** Higher order interactions, species coexistence, modern coexistence theory, pairwise coexistence

## Abstract

Higher order interactions (HOIs) have been suggested to stabilize diverse ecological communities. However, their role in maintaining species coexistence from the perspective of modern coexistence theory is unknown. Here, using a three-species Lotka-Volterra model, we derive a general rule for species coexistence modulated by HOIs. We show that negative HOIs that intensify pairwise competition, can promote coexistence across a wide range of fitness differences, provided that HOIs strengthen intraspecific competition more than interspecific competition. In contrast, positive HOIs that alleviate pairwise competition can also stabilize coexistence across a wide range of fitness differences, irrespective of differences in strength of inter- and intraspecific competition. Furthermore, we extend our three-species analytical result to multispecies competitive community and show, using simulations, that feasible multispecies coexistence is possible provided that strength of negative intraspecific HOIs is higher than interspecific HOIs. In addition, multispecies communities, however, become unstable with positive HOIs as such higher-order interactions could lead to disproportionately infeasible growth rates. This work provides crucial insights on the underlying mechanisms that could maintain species diversity and links HOIs with modern coexistence theory.

## 1. Introduction

In diverse ecological communities, species interact with other species in the community temporally (Li & Chesson, 2016) and/or spatially (Hart *et al.*, 2017). Understanding the mechanisms behind maintenance of species diversity has been one of the central goals of ecological research for decades (Levine *et al.*, 2017). While our primary understanding of species coexistence empirically and theoretically has mostly come from a study of species pairs (Chesson, 2000; Hart *et al.*, 2016; Kraft *et al.*, 2015), understanding the dynamics and coexistence of multiple species from the viewpoint of species pairs becomes unfeasible and intractable as species richness increases (Barabas *et al.*, 2016; Bairey *et al.*, 2016).

In competition models, the underlying processes that facilitate species coexistence require parameter trade-offs in competitive interactions to stabilize multispecies coexistence (Barabas *et al.*, 2016). Moreover, competitive interactions between species in a diverse community is difficult to structure based on the generalizations coming from simple coexistence rules, leading to formulation of models of coexistence based on ecological equivalence (Hubbell, 2006). In species-rich communities, underlying processes that cannot be captured by pairwise species interactions can emerge, and have been suggested to promote species coexistence(Abrams 1983). Such underlying processes, for example can be intransitive or “rock-paper-scissors” interactions (Laird & Schamp, 2006; Gallien *et al.*, 2017; Saavedra *et al.*, 2017). Intransitive interactions are inherently pairwise in nature but they form interaction chains that favor species coexistence. For example, in a three species system, intransitive competitive interactions can lead to coexistence of all the three species, although none of the species pair can coexist alone. Advancing our understanding of species coexistence in diverse communities requires an understanding of mechanisms that could inherently be high-dimensional.

When diverse species communities are modelled explicitly by considering only pairwise interactions, multiple species coexistence is not always possible (Levine *et al.*, 2017; Bairey *et al.*, 2016; Wilson, 1992). Recent studies have suggested that in order for large number of species to coexist, interactions between species should not be constrained to species pairs but should include higher order interactions (HOIs) (Grilli *et al.*, 2017). Although, such HOIs have been suggested to stabilize large ecological communities, their underlying role in maintaining species coexistence, from the purview of modern coexistence theory (MCT) is unknown. This is primarily due to the difficulty of integrating MCT for more than two species simultaneously (Saavedra *et al.*, 2017; Barabas *et al.*, 2016). MCT states that coexistence is possible when fitness differences between species are smaller than their niche differences (Chesson, 2000). In MCT, coexistence of species can be understood from a mutual invasibility criteria, where the invasion growth rate of a species is analytically decomposed into stabilizing niche differences and average fitness differences (Kremer & Klausmeier, 2013; Gallien *et al.*, 2017). Niche differences increase the probability for species coexistence while fitness differences increase the probability of competitive exclusion. Importantly, fitness and niche differences can be quantified from the terms of the Lotka-Volterra pairwise competition model as (Saavedra *et al.*, 2017):

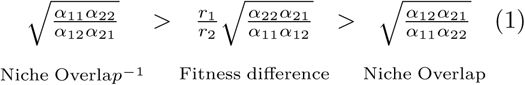

If niche overlap is greater than the fitness difference between two species, then coexistence is not possible. Under certain simplistic assumptions, one can integrate results from HOIs into the traditional framework of pairwise species coexistence. Such an integration would make the relevance and the understanding of HOIs more complete.

Here, using a three-species Lotka-Volterra model, we demonstrate the importance of HOIs in maintaining and disrupting species coexistence. Specifically, using invasibility criterion, we modified pairwise interspecific coefficients of the Lotka-Volterra model in a way that allowed us to create a range of fitness differences ranging from low to high. We then show how negative three-way HOIs (HOIs that intensify pairwise competitions), and positive three-way HOIs (HOIs that alleviate pairwise competition) can stabilize species coexistence in fitness regions, where species coexistence is impossible if only pairwise interactions are considered. We then extend our three-species HOIs case, to a multispecies competitive community, and show that the conditions under which HOIs stabilize species coexistence in the three-species case still holds in a multispecies community. We highlight the possible mechanisms by which HOIs could promote coexistence in species-rich communities.

## 2. Methods

### 2.1 Higher order interactions

Lotka-Volterra model for three-way HOIs can be written as (Letten and Stouffer 2019b) (for four-way HOIs see appendix A):

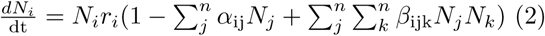

where *α*_ij_ and *β*_ijk_ are pair-wise interactions and HOIs respectively. Here, higher-order terms could broadly be defined as non-additive effects on per capita fitness of a species. HOIs could intensify or alleviate the pairwise competition between two species depending on the sign of *β*_ijk_ as negative or positive respectively. Here, *n* = 3, in this particular section, and we evaluate the effect of HOIs for multispecies (n = 50) communities later (see section 2.3). In this particular model, we make a few assumptions while deriving the invasion growth rate for the three species case-

1) There is interspecific competitive interaction between species 1 and 2, but not with species 3. This means that the matrix of competitive interactions will be:

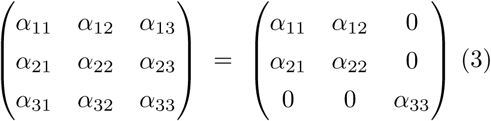

2) Only interspecific HOIs are taken into account. This means that terms such as *β*_iii_ = 0, where *i* = 1, 2, 3.

3) Species 3 influences species 1, and species 2 only through HOIs. However, species 3 does not get influenced by species 1 or 2 through HOIs (*β*_3*jk*_ = 0; where *j, k* = 1, 2).

These assumptions were made to ensure that the number of terms in the HOI model are tractable for simple analysis (appendix A). Using these assumptions, we can expand the model in equation (2) as:

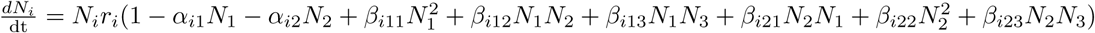

where *i* = 1, 2 (for species 3 see appendix A). We can then write the HOI matrix for each of the species as:

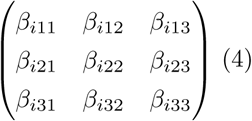

Where *i* = 1, 2; *β*_*i*31_ = *β*_*i*32_ = *β*_*i*33_ = 0; and *β*_iii_ = 0.

We consider cases of both positive and negative HOIs, while calculating the invasion growth rates.

### 2.2 Invasion growth rate and coexistence theory

The invasion growth rate, *r*_*i*_, is the per capita rate of increase in a species’ abundance—when it is rare—in presence of the other species, which is at equilibrium in the community. This means 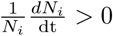 for species *i* in the community. Invasion growth rates of species 1 in the presence of HOIs in the community can be written as (see appendix A):

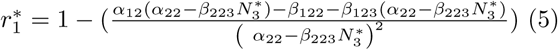

where 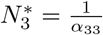 (appendix A).

We evaluated the effect of 3-way and 4-way HOIs in promoting coexistence by comparing invasion growth rates of species 1 in presence and absence of HOIs. Following Gallien et. al 2017, we created scenarios where pairwise competitive matrix (3) varied from purely symmetric pairwise interactions to asymmetric interactions with gradually increasing pairwise fitness differences. As pairwise fitness differences (calculated from equation 1) increase, niche difference (calculated from equation 1) should increase accordingly to stabilize species coexistence. This was done by modifying the pairwise interaction matrix to (Gallien *et al.*, 2017):

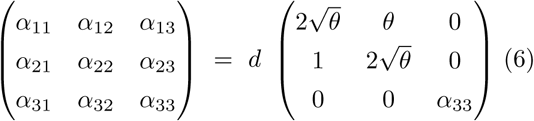

Where d = 0.01. The above modified matrix ensures that niche overlap between species 1 and species 2 is at 0.5 even when fitness differences (calculated using equations 1), controlled by *θ*, increases linearly. As *θ* is varied from 0 to 7, fitness difference between species 2 and 1 increases linearly. Note that fitness difference of species 2 over species 1 is given from equation 1 as 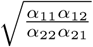. For certain *θ* = [5, 7] values, fitness difference of species 2 over species 1 exceeds niche difference. Consequently, following the pairwise coexistence rule, species 1 can never invade species 2. This means that pairwise coexistence is impossible for certain *θ* = [5, 7] values. *θ* = 5 defines the boundary between stable pairwise coexistence and competitive exclusion, when there are no HOIs. Note that species 3 remains unaffected by this modification and only participates in HOIs.

Next, to evaluate whether the presence of three-way and four-way HOIs (appendix B) stabilizes pairwise coexistence in scenarios where fitness differences are extreme, we estimated invasion growth rates of species 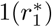 when species 2 is present at equilibrium. When HOIs are absent, it is expected that given niche overlap is at 0.5, pairwise coexistence becomes impossible as fitness differences increase. The importance of HOIs will be evident if species 1 could increase its invasion growth rates and invade when HOIs are present even when fitness differences are large, i.e. they could increase their numbers even when differences between the two species in terms of fitness are large. For sensitivity analysis of invasion growth rate to HOIs see appendix C.

### 2.3 Multispecies coexistence and higher order interactions

Multispecies generalization of two-species pairwise coexistence rule, however, is complicated. In multispecies communities, while all species pairs must satisfy the pairwise coexistence rule, this does not, however, guarantee stability (Barabas *et al.*, 2016). For multispecies competitive communities, the stability and feasibility of coexistence can be evaluated from Weyl’s inequality (see below). A pairwise competitive community can be written as:

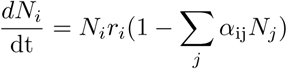

Where, *α*_ij_ represents pairwise competitive interactions. *α*_ij_ is the element in the *i-th* row and *j-th* column of a matrix of pairwise competitive interactions. This pairwise matrix of competitive interactions can be denoted, by, say **A**. Here, we consider **A** as a symmetric matrix, such that, *α*_ij_ = *α*_ji_.

Now, **A** can be decomposed into inter- and intraspecific matrices, say **B** and **C** respectively, where **B** is a matrix of only interspecific competitive interactions while **C** is a matrix of intraspecific competitive interactions. Now, since **C** contains only intraspecific coefficients, **C** is a diagonal matrix, whereas **B** has zeros in the diagonal. The off-diagonal entries of **B** capture the symmetric pairwise interspecific competitive interactions. Hence, we can write, **A = B + C.** Since, **A** is symmetric, **B** and **C** are also symmetric, and hence all their eigenvalues are real. For a *S* species community, **B**, and **C** will have *S* eigenvalues and these eigenvalues can be ordered as *b*_1_ ≥ *b*_2_ ≥ *b*_3_ ≥ *…* ≥ *b*_*s*_ and *c*_1_ ≥ *c*_2_ ≥ *c*_3_ ≥ *…* ≥ *c*_*s*_. The Weyl’s inequality now states that the necessary and sufficient condition for multispecies coexistence to be stable is:

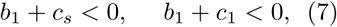

When all intraspecific coefficients (or the diagonal elements of **C)** are equal, all the eigenvalues have the same value i.e., *c*_*s*_ = *c*_1_ = *c*, and the necessary and sufficient condition for stable species coexistence then becomes (Barabas *et al.*, 2016):

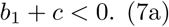

We thus structured our analysis in the following way – we took a 50 species competitive community where intraspecific competition coefficients, i.e., diagonal elements of **C** were kept the same. The interspecific competitive interactions of **B** were drawn from a random uniform distribution in a way that it either satisfied the Weyl’s inequality or it did not. Typically, Weyl’s inequality was fulfilled when intraspecific effects were much larger than all the interspecific effects (Barabas *et al.*, 2016); and the inequality of (7a) was not satisfied whenever random (but symmetric) interspecific interactions drawn from a uniform distribution had similar or higher values than intraspecific effects. When pairwise competitive interactions satisfied Weyl’s inequality (7a), coexistence of all 50 species was stable (Fig. 3C), and when it did not, coexistence was unstable (Fig. 3D).

**Figure 1:**
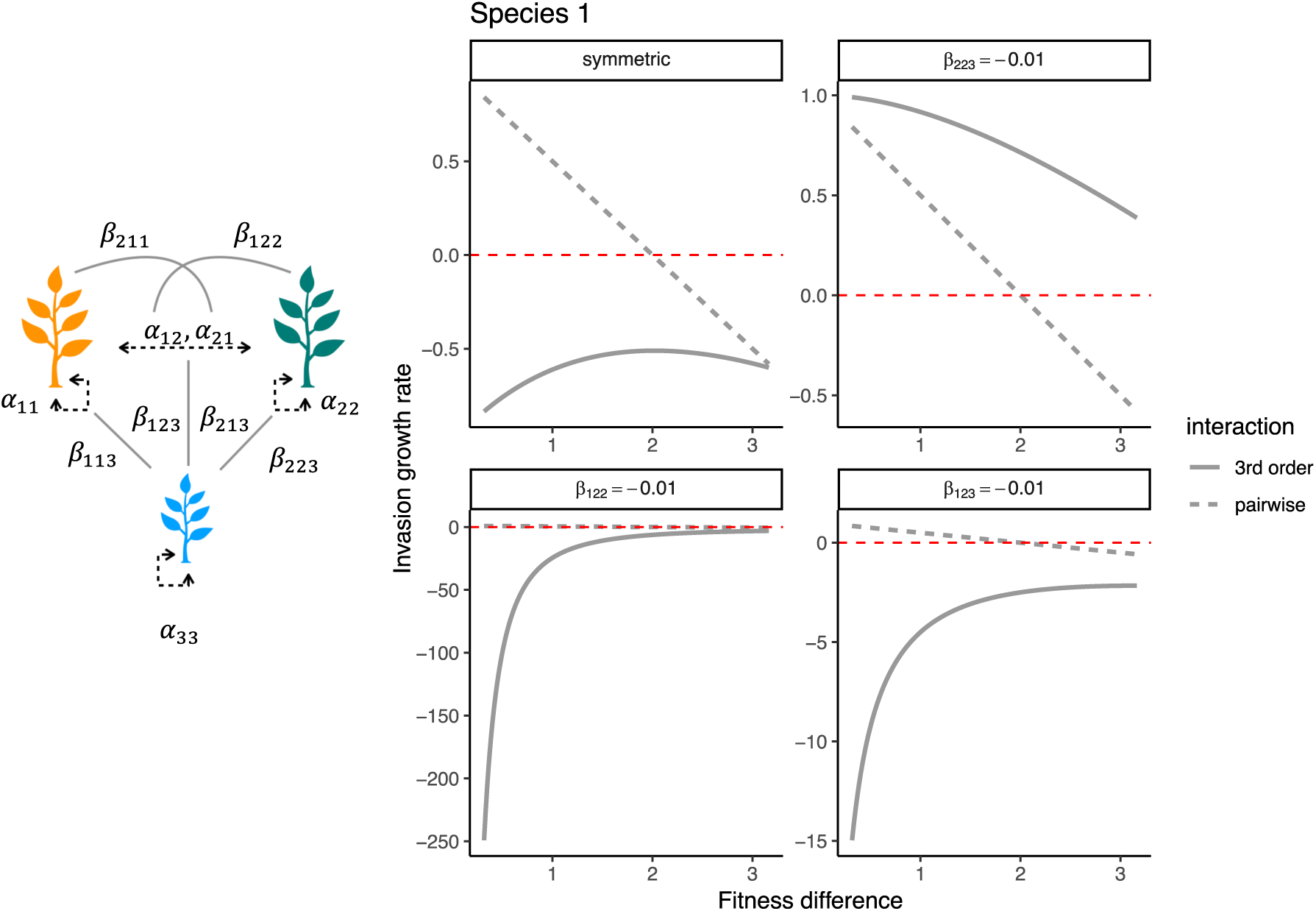
Invasion growth rate (Y-axis) of species 1 (top row) for pairwise species competition (dashed grey lines) and negative three-way HOIs (grey solid lines) for a range of fitness difference (X-axis). The red-dashed line marks the y-intercept at zero. Each panel of the plot compares invasion growth rate of species 1 under pairwise competition and under negative three-way HOIs, with the panel label referring to the values of the HOI terms. For instance, the panel “symmetric” would mean that all the HOI terms in the HOI matrix have the exact same magnitude of -0.01; and the panel *β*_223_= -0.01 (top row, species 1) would mean all the elements of HOI matrix are zero except *β*_223_ which is at -0.01 (i.e., more negative would mean more increase in strength of intraspecific competition of species 2, *α*_22_). Panels *β*_122_= -0.01 and *β*_123_= -0.01, would mean all terms of HOI matrix are zero except *β*_122_ and *β*_123_ respectively. Note that invasion growth rate of species 1 (top row) was negative across the range of fitness difference in negative three-way HOIs when interspecific competition was intensified more than intraspecific competition (panels: *β*_122_ and *β*_123_ for species 1).

**Figure 2:**
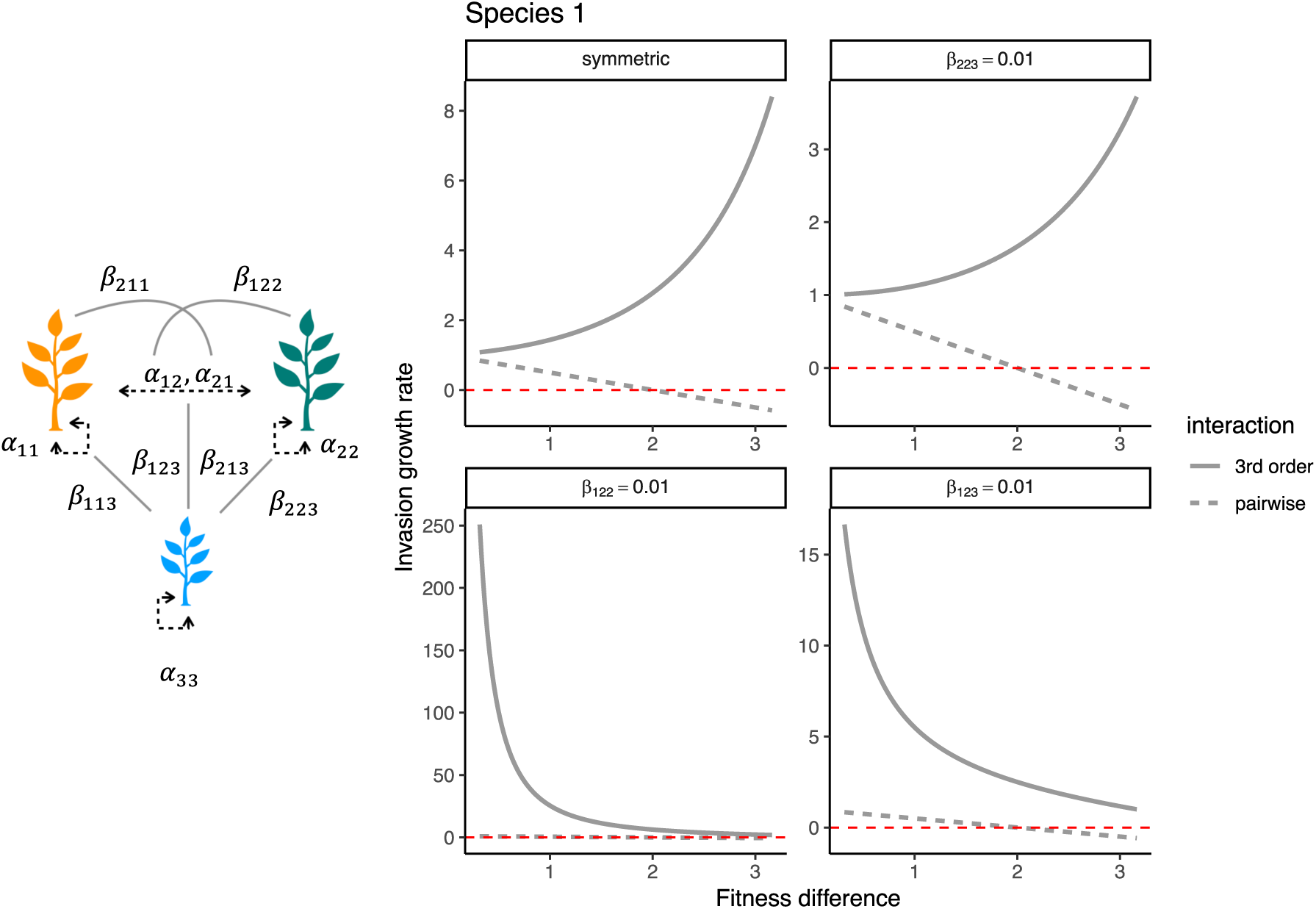
Invasion growth rate (Y-axis) of species 1 (top row) for pairwise species competition (dashed grey lines) and positive three-way HOIs (grey solid lines in the figure) shown for a range of fitness difference (X-axis). The red-dashed line marks the y-axis at zero invasion growth rate. The panel label refers to the values of the terms in the HOI matrix. For instance, the panel “symmetric” would mean that all the HOI terms in the HOI matrix have the exact same magnitude of 0.01; and the panel *β*_223_= 0.01 (top row, species 1) would mean all the elements of HOI matrix are zero except *β*_223_ which is at 0.01 (i.e., more positive and hence more decrease in strength of intraspecific competition of species 2, *α*_22_). Panel *β*_122_= -0.01 and panel *β*_123_ = 0.01, would mean all terms of HOI matrix are zero except *β*_122_ and *β*_123_ respectively. Note that invasion growth rate of species 1 (top row) was positive across the range of fitness difference in positive three-way HOIs (panels: *β*_122_ and *β*_123_ for species 1).

**Figure 3:**
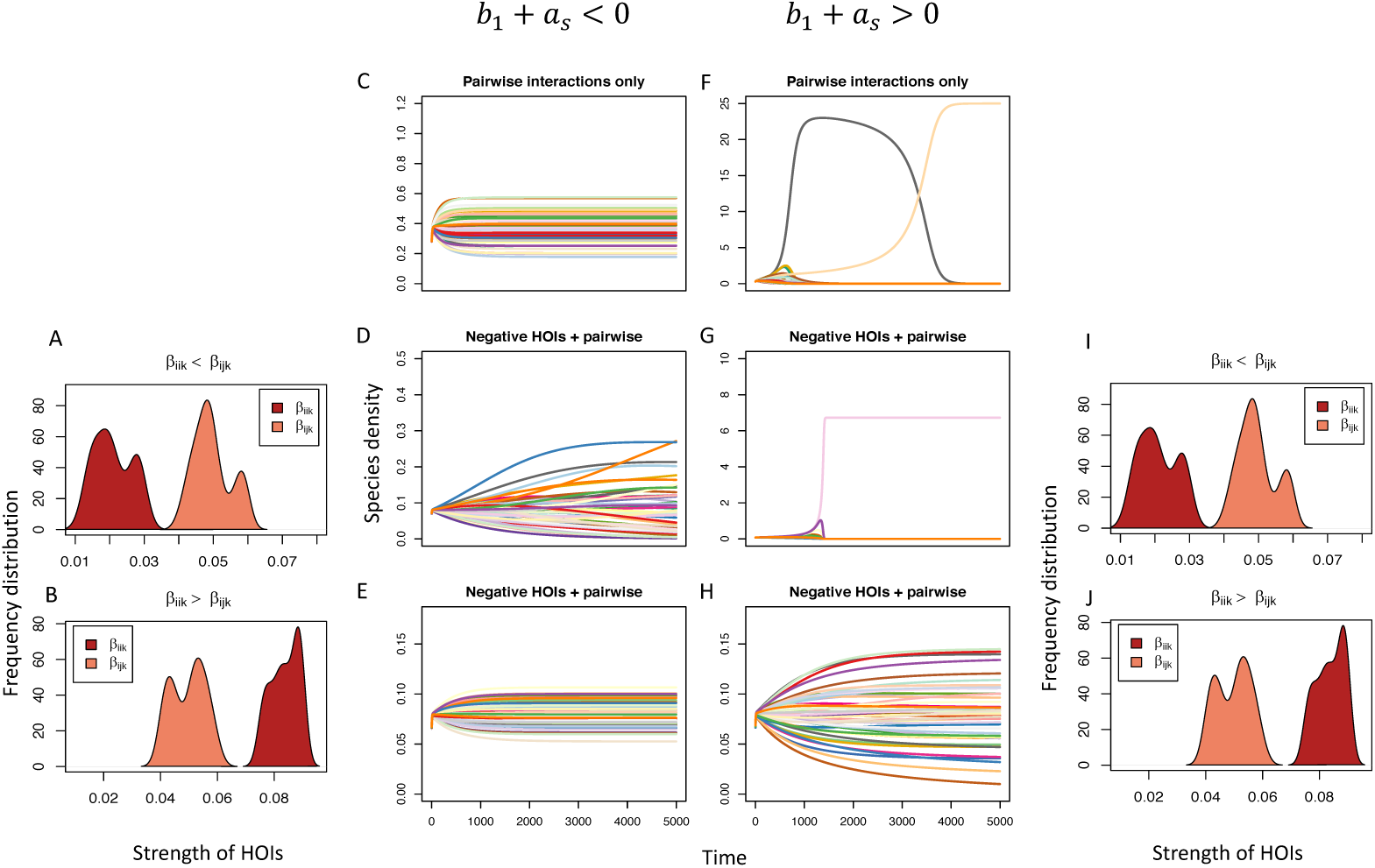
Multispecies coexistence in the presence and absence of negative HOIs that either satisfy Weyl’s inequality (b_1_+a_s_ < 0) or do not (b_1_+a_s_ > 0). When 50 species in a competitive community compete in pairwise manner, and satisfies Weyl’s inequality (C) species coexistence was always stabilized. However, in the presence of negative HOIs where interspecific HOIs were strictly stronger than intraspecific HOIs, i.e., *β*_ijk_ *> β*_iik_ (A), species coexistence was destabilized (D), while when the opposite happens, i.e., *β*_iik_ *> β*_ijk_ (B), which suggests intraspecific HOIs to be stronger than interspecific HOIs, species coexistence was again stabilized (E). When Weyl’s inequality was not satisfied, (b_1_+a_s_ > 0) (F), pairwise coexistence was impossible (F). However, in the presence of negative HOISs and if intraspecific HOIs were stronger than interspecific HOIs, *β*_iik_ *> β*_ijk_ (J), species coexistence was then stabilized (H), but disrupted again (G) if interspecific HOIs were stronger than intraspecific HOIs (I).

Next, we wanted to evaluate the circumstances under which negative and positive HOIs could stabilize multispecies coexistence, when pairwise competitive interactions did not satisfy Weyl’s inequality *b*_1_ + *c*_1_ ≮ 0 (i.e. when pairwise species coexistence is destabilized). We assembled interspecific and intraspecific HOIs from random uniform distributions and investigated the effects on multispecies coexistence. In addition, we wanted to investigate whether HOIs can destabilize species coexistence when Weyl’s inequality is fulfilled (i.e. species coexistence is stabilized by pairwise interactions).

## 3. Results

### 3.1 Negative higher-order interactions

In the simple pairwise interaction case, given a niche overlap of 0.5, coexistence between species 1 and 2 was only possible when fitness differences between them ranged from 0 to 2. Beyond a fitness difference of 2, coexistence was not possible, as the invasion growth rate became negative.

Interestingly, we found that, when three-way HOIs were negative, invasion growth rate of species 1 was positive across the range of fitness differences, provided species 3 intensified intraspecific competition of species 2 (*β*_223_) more than it intensified interspecific competition (*β*_123_) (Fig. 1). However, if all negative three-way HOIs had the same magnitude, species coexistence was impossible even with low fitness differences (Fig. 1, symmetric case).

Negative four-way HOIs could also promote coexistence, even when fitness differences between two species were high, if their strength was an order of magnitude lower in comparison to inter and intraspecific pairwise competition and three-way higher order interactions (appendix B, Fig. B1-2, B5-6).

### 3.2 Positive higher-order interactions

Generally, positive three-way HOIs could lead to species coexistence despite substantial fitness difference between the two species and even in fitness regions where species coexistence is impossible if only pairwise interactions are considered (Fig. 2). For example, when species 3 alleviated intraspecific competition of species 2 (*β*_223_) more than it alleviated interspecific competition (of species 2 on 1), invasion growth rate of species 1 increased non-linearly, as fitness difference increased. This particular result could be understood by looking at the invasion growth rate of species 1 in the presence of non-zero *β*_223_ HOI (i.e. effect of species 3 on intraspecific interaction of species 2) while rest of the HOI terms are zero), which becomes:

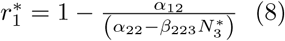

As fitness difference increased, due to *θ* varying from 0 to 7 in (3), interspecific effect of species 2 on species 1 increased more rapidly than intraspecific competition of species 2. Because as *θ* increases in (6), interspecific effects *α*_12_ increases at the same rate as *θ*, while intraspecific effects *α*_22_ in (6) increases by 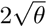. With 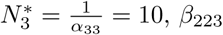 of 0.01 causes the invasion growth rate of species 1 (8) to increase rapidly as fitness difference increases. In all the cases, positive HOIs lead to high invasion growth rates.

Positive four-way HOIs led to species coexistence despite extreme fitness differences. As strength of positive four-way HOIs increased, invasion growth rate of species 1 when species 2 and species 3 are present, also increased (appendix Fig. B7-8).

### 3.3 Higher-order interactions and coexistence in a large competitive community

When Weyl’s inequality was satisfied by pairwise competitive interactions, a 50 species community was feasible and stable in the absence of HOIs (Fig. 3 C). Failing to fulfil the inequality led to disruption of pairwise species coexistence (Fig. 3G).

Interestingly, in the presence of negative HOIs, even when Weyl’s inequality was not fulfilled, coexistence of 50 species was possible provided intraspecific HOIs were stronger than interspecific HOIs (Fig. 3H, 3J). When interspecific HOIs were stronger than intraspecific HOIs, coexistence of all 50 species was impossible, irrespective of whether Weyl’s criteria was satisfied or not (Fig. 3D, 3A). Thus, failing to satisfy Weyl’s criteria, stronger interspecific HOIs than intraspecific HOIs compounded the disruption of species coexistence.

Positive three-way HOIs led to unfeasible species densities when their magnitude was of similar strength to that of pairwise interactions. However, under certain parameter choices (low values of HOIs) of positive HOIs strength, coexistence of all 50 species was possible despite differences in strength of intraspecific or interspecific HOIs, provided Weyl’s inequality was satisfied.

## 4. Discussion

While the assumption that interactions between species in a species rich community is inherently pairwise is pervasive in coexistence theory (Gallien *et al.*, 2017; Terhorst *et al.*; Levine *et al.*, 2017). The role of HOIs in stabilizing or destabilizing coexistence has been relatively understudied both empirically and theoretically (Letten & Stouffer, 2019; Grilli *et al.*, 2017; Baruah & John, 2019; Abrams, 1983). Lately, empirical understanding of species coexistence has been sought through modern coexistence theory where invading potential of a species in the presence of an established competitor species is explored (Grainger *et al.*, 2019). In this context, HOIs need to be elucidated clearly to fully understand multispecies coexistence, and the underlying mechanisms that leads to species coexistence (Levine *et al.*, 2017). Our study fills this gap by using concepts from modern coexistence theory (Chesson, 2000) to evaluate the effect of HOIs on species coexistence in a simple three species Lotka-Volterra model and in a more complex multispecies community. This study, in general, shows that HOIs promote species coexistence in parameter spaces where pairwise coexistence is unstable, provided certain conditions are fulfilled.

Using a three species Lotka-Volterra model, our results showed that positive HOI’s stabilize coexistence across a wide range of fitness differences irrespective of differences in strength of inter- and intraspecific competition, while negative HOI’s stabilize coexistence only if intraspecific competition was strengthened more than interspecific competition. If, however, in the case of negative HOIs, interspecific competition was strengthened more in relation to intraspecific competition (intraspecific HOIs < interspecific HOIs), species coexistence was impossible even when fitness differences were negligible. This particular result has been observed in another eco-evolutionary study, where negative density-mediated HOIs have been shown to promote multispecies coexistence (Baruah & John, 2019).

Positive HOIs lead to a decrease in the per-capita strength of competition between two species, while negative HOIs led to an increase in the per-capita strength of competition in our modelling scenario (Bairey *et al.*, 2016). Moreover, positive HOIs could lead to disproportionately high invasion growth rates, particularly when it alleviated intraspecific competition. Such high invasion growth rate could affect stability and lead to infeasible species densities (Terry *et al.*, 2018)(appendix Fig. C13). Importantly, when fitness differences between two species were extremely high, we believe positive HOIs could lead to species coexistence by decreasing interspecific competition more than intraspecific competition. For example, for extreme fitness difference between species 1 and species 2, say fitness difference of 3, positive HOIs (*β*_123_ = 0.01), which decreased interspecific competition of species 2 on species 1, will lead to species coexistence and proportionately low but positive invasion growth rate (Fig. 2). An earlier study had reported that HOIs could positively influence fitness of species by alleviating the dominating effect of neighboring species (Bairey *et al.*, 2016; Mayfield & Stouffer, 2017).

Our results suggested that invasion growth rates were generally sensitive to changes in the strength of HOIs, for both positive and negative HOIs (appendix Fig. C9), which suggests that parameter changes in HOIs has the potential to destabilize species coexistence. Hence under restricted parameter space, HOIs could stabilize species coexistence (AlAdwani & Saavedra, 2019). Four-way HOIs could lead to species coexistence across a range of fitness differences, if their direction was positive (appendix Fig. B3-4; B7-8). Negative four-way HOIs could also promote coexistence, provided their strength was an order of magnitude lower in comparison to pairwise interactions (appendix Fig. B1-2, B5-6). It is possible that four-way interactions could be prevalent in species communities, but empirically parameterizing such four-way interactions would be a difficult task (although see Mayfield & Stouffer (2017)). In addition, incorporating four-way HOIs in models parameterized from empirical data might not always provide additional explanatory power (AlAdwani & Saavedra, 2019).

In a multi-species community, pairwise coexistence rule does not hold (Barabas *et al.*, 2016; Saavedra *et al.*, 2017). However, multispecies coexistence could be understood by analyzing the Weyl’s inequality (Fulton, 2000). When Weyl’s inequality was not satisfied, pairwise species coexistence was impossible. In our models, positive HOIs decreased pairwise competition in a multispecies community, which consequently led to species coexistence provided self-regulation was strong (Mayfield & Stouffer, 2017) and Weyl’s inequality was satisfied (appendix Fig. C13, column *b*_1_ +*c*_1_ < 0). When Weyl’s equality was not satisfied, species coexistence was destabilized both with pairwise and with positive HOIs. Pairwise coexistence in such a scenario was disrupted particularly because strength of intraspecific competition was on average lower than interspecific competition between species. However, surprisingly, even in such a scenario, negative HOIs could still stabilize species coexistence. Particularly, if intraspecific HOIs was strictly greater than interspecific HOIs, species coexistence in a large competitive community was stabilized. This is analogous to the two-species coexistence rule, that species must limit themselves more than they limit competitors. In general, the simplest way to generalize multispecies coexistence in the presence of negative HOIs was that – when pairwise coexistence for multispecies community was impossible (Weyl’s inequality not satisfied), intraspecific competition should be strengthened more than interspecific competition by HOIs. If even a single species in the multispecies community violated this rule, coexistence in the multispecies community was disrupted, even in the presence of HOIs (appendix Fig. C11). On the contrary, when pairwise multispecies coexistence was possible (Weyl’s inequality was satisfied and pairwise intraspecific effects were substantially larger than interspecific effects), having intraspecific HOIs that had similar strength as interspecific HOIs could still stabilize species coexistence (appendix Fig. C12). This particular result was also reiterated by another study that dealt with eco-evolutionary processes that might emerge when species in a large competitive community had density-mediated HOIs (Baruah & John, 2019). The study showed that when negative HOIs were present, species not only coexisted but their traits also evolved to be very similar. Moreover, species structured themselves in a trait axis over evolutionary time in a way that negative HOIs strengthened pairwise intraspecific competition more than it intensified interspecific competition (Baruah & John, 2019). Furthermore, from our simple three-species results, we observed that positive HOIs could lead to disproportionately high invasion growth rates and henceforth might negatively affect stability (Terry *et al.*, 2018, 2017).

Lotka-Volterra models of competition have been extensively used to understand mechanisms that could promote species coexistence through pairwise interactions (Barabas *et al.*, 2016; Hart *et al.*, 2016) and through HOIs (Wilson, 1992; Bairey *et al.*, 2016; Letten & Stouffer, 2019). Using Lotka-Volterra models, our results add to this emerging body of knowledge on HOIs and the prospect of extending simple tractable dynamics to understanding complex multispecies dynamics (Wilson, 1992; Bairey *et al.*, 2016; Letten & Stouffer, 2019; Baruah & John, 2019; Grilli *et al.*, 2017; Terry *et al.*, 2017). It is straightforward, although challenging, to estimate pairwise competition coefficients from experiments that involve manipulating competitor densities, or from long term observational field data. However, there are other obstacles in estimating higher-order coefficients and in understanding heir effects on species coexistence (Mayfield & Stouffer, 2017). For instance, to implement a higher-order model empirically with just three competitors would require no less than 27 parameters at the very least. To collect such amount of empirical data whether observational or experimental is an enormous task. Nonetheless, invasion growth rates could be estimated from empirical data that are used to parameterize community models or mechanistic competition models by explicitly incorporating HOIs (Tilman, 1994; Letten & Stouffer, 2019).

Although we have much to understand about the effects of HOIs empirically, it is clear that effects of HOIs on species coexistence is dependent on their strength as well as on their direction. That being said, HOI terms in ecological models could increase the number of equilibrium points exponentially, though such equilibrium points can be ecologically feasible only under a restricted set of parameter space (AlAdwani & Saavedra, 2019).

## Acknowledgements

The authors have no conflicts of interests and would like to thank Jordi Bascompte for discussions on an earlier version of the manuscript. This research was funded by Forschungskredit UZH FK-18-082 to GB.

## Data availability

Data will be made available through the Dryad repositories.

## References

Abrams, P.A. (1983) Arguments in Favor of Higher Order Interactions. The American Naturalist 121, 887–891.

AlAdwani, M. & Saavedra, S. (2019) Is the addition of higher-order interactions in ecological models increasing the understanding of ecological dynamics? Mathematical Biosciences 315, 108222.

Bairey, E., Kelsic, E.D. & Kishony, R. (2016) High-order species interactions shape ecosystem diversity. Nature Communications 7, 12285–12285.

Barabas, G., D’Andrea, R. & Vasseur, D. (2016) The effect of intraspecific variation and heritability on community pattern and robustness, vol. 19. Blackwell Publishing Ltd.

Baruah, G. & John, R. (2019) Intraspecific variation promotes species coexistence and trait clustering through higher order interactions. bioRxiv p. 494757.

Chesson, P. (2000) Mechanisms of Maintenance of Species Diversity. Annual Review of Ecology and Systematics 31, 343–366.

Fulton, W. (2000) Eigenvalues, invariant factors, highest weights, and Schubert calculus. Bulletin of the American Mathematical Society 37, 209–249.

Gallien, L., Zimmermann, N.E., Levine, J.M. & Adler, P.B. (2017) The effects of intransitive competition on coexistence. Ecology Letters 20, 791–800.

Grainger, T.N., Levine, J.M. & Gilbert, B. (2019) The Invasion Criterion: A Common Currency for Ecological Research. Trends in Ecology & Evolution.

Grilli, J., Barabás, G., Michalska-Smith, M.J. & Allesina, S. (2017) Higher-order interactions stabilize dynamics in competitive network models. Nature 548, 210–210.

Hart, S.P., Schreiber, S.J., Levine, J.M. & Coulson, T. (2016) How variation between individuals affects species coexistence. Ecology Letters 19, 825–838.

Hart, S.P., Usinowicz, J. & Levine, J.M. (2017) The spatial scales of species coexistence. Nature Ecology & Evolution 1, 1066–1073.

Hubbell, S.P. (2006) Neutral Theory and the Evolution of Ecological Equivalence. Ecology 87, 1387–1398.

Kraft, N.J.B., Godoy, O. & Levine, J.M. (2015) Plant functional traits and the multidimensional nature of species coexistence. Proceedings of the National Academy of Sciences of the United States of America 112, 797–802.

Kremer, C.T. & Klausmeier, C.A. (2013) Coexistence in a variable environment: Eco-evolutionary perspectives. Journal of Theoretical Biology 339, 14–25.

Laird, R.A. & Schamp, B.S. (2006) Competitive intransitivity promotes species coexistence. The American naturalist 168, 182–93.

Letten, A.D. & Stouffer, D.B. (2019) The mechanistic basis for higher-order interactions and nonadditivity in competitive communities. Ecology Letters 22, 423–436.

Levine, J.M., Bascompte, J., Adler, P.B. & Allesina, S. (2017) Beyond pairwise mechanisms of species coexistence in complex communities. Nature 546, 56–64.

Li, L. & Chesson, P. (2016) The Effects of Dynamical Rates on Species Coexistence in a Variable Environment: The Paradox of the Plankton Revisited. The American naturalist 188, E46–58.

Mayfield, M.M. & Stouffer, D.B. (2017) Higher-order interactions capture unexplained complexity in diverse communities. Nature Ecology & Evolution 1, 0062–0062.

Saavedra, S., Rohr, R.P., Bascompte, J., Godoy, O., Kraft, N.J.B. & Levine, J.M. (2017) A structural approach for understanding multispecies coexistence. Ecological Monographs 87, 470–486.

Terhorst, C.P., Zee, P.C., Heath, K.D., Miller, T.E., Pastore, A.I., Patel, S., Schreiber, S.J., Wade, M.J. & Walsh, M.R. (????) Evolution in a Community Context: Trait Responses to Multiple Species Interactions*.

Terry, J.C.D., Morris, R.J. & Bonsall, M.B. (2017) Trophic interaction modifications: an empirical and theoretical framework. Ecology Letters 20, 1219–1230.

Terry, J.C.D., Morris, R.J. & Bonsall, M.B. (2018) Trophic interaction modifications disrupt the structure and stability of food webs. bioRxiv p. 345280.

Tilman, D. (1994) Competition and biodiversity in spatially structured habitats. Ecology 75, 2–16.

Wilson, D.S. (1992) Complex Interactions in Metacommunities, with Implications for Biodiversity and Higher Levels of Selection. Ecology 73, 1984–2000.

