## appendix for "Higher order interactions and coexistence theory"

---

*Keywords:* higher order interactions, modern coexistence theory

---

### Appendix A. Three-way higher order Lotka-Volterra model and invasion growth rates

$$\frac{dN_i}{dt} = N_i r_i (1 - \sum_j \alpha_{ij} N_j + \sum_j \sum_k \beta_{ijk} N_j N_k) \tag{A.1}$$

Here, $N_i$ refers to density of species $i$, $\alpha_{ij}$ and $\alpha_{ijk}$ are the pair-wise and higher order interactions [1, 2, 4]. Above equation can be expanded to for species 1 :

$$\frac{dN_1}{dt} = N_1 r_1 (1 - \alpha_{11} N_1 - \alpha_{12} N_2 + \beta_{112} N_1 N_2 + \beta_{113} N_1 N_3 + \beta_{121} N_2 N_1 + \beta_{122} N_2^2 + \beta_{123} N_2 N_3) \tag{A.2}$$

And for species 2 as:

$$\frac{dN_2}{dt} = N_2 r_2 (1 - \alpha_{21} N_1 - \alpha_{22} N_2 + \beta_{212} N_1 N_2 + \beta_{213} N_1 N_3 + \beta_{221} N_2 N_1 + \beta_{211} N_1^2 + \beta_{223} N_2 N_3) \tag{A.3}$$

Species 3 however not compete with species 1 and 2, but instead influences pairwise competition between species 1 and 2 through higher order interactions. For instance, species 3, could be a functionally different plant species. Being functionally different allows species 3 to occupy a different niche and thus has no direct competitive affect on either species 1 or species 2. Because it is functionally different, it could be possible that it alters its environment that indirectly affects pairwise competition between species 1 and 2. For example, species 3 could alter the soil rhizosphere that ultimately impacts competitive interactions of species 1 and 2 [3, 5]. Species 3 dynamics can be written as :

$$\frac{dN_3}{dt} = N_3 r_3 (1 - \alpha_{33} N_3) \tag{A.4}$$

Isoclines of $N_1$ and $N_2$ can be found from equation 3 and 4 as :

sc

$$N_1 = \frac{r_2 - \alpha_{12} N_2 + \beta_{123} N_2 N_3 + \beta_{122} N_2^2}{\alpha_{11} - \beta_{112} N_1 - \beta_{113} N_3 - \beta_{121} N_2}$$

and,

$$N_2 = \frac{r_1 - \alpha_{21} N_1 + \beta_{213} N_1 N_3 + \beta_{211} N_1^2}{\alpha_{22} - \beta_{212} N_1 - \beta_{223} N_3 - \beta_{221} N_1}$$

With this, we can calculate the invasion growth rate of species 1. In simple terminology, invasion growth rate of a species is the rate of increase in its density when rare. Invasion growth rate of a species depends on the invasibility criteria that determines the conditions for which $\frac{dN_1}{dt}\frac{1}{N_1} > 0$. Invasion growth rate of species 1 can thus be calculated by :

$$\frac{dN_1}{dt}\frac{1}{N_1} > 0,$$

which leads to,

$$(1 - \alpha_{11} N_1 - \alpha_{12} N_2 + \beta_{112} N_1 N_2 + \beta_{113} N_1 N_3 + \beta_{121} N_2 N_1 + \beta_{122} N_2^2 + \beta_{123} N_2 N_3) > 0,$$

Species 1, when rare ($N_1 = 0$) can invade species 2 and species 3 when they are at equilibrium, if

$$(1 - \alpha_{12} N_2 + \beta_{122} N_2^2 + \beta_{123} N_2 N_3) > 0, \tag{A.5}$$

which leads to after substituting $N_2$'s isocline ,

$$r_1^* = 1 - \Big(\frac{\alpha_{12}(\alpha_{22} - \beta_{223}N_3^*) - \beta_{122} - \beta_{123}(\alpha_{22} - \beta_{223}N_3^*)}{(\alpha_{22} - \beta_{223}N_3^*)^2}\Big). \tag{A.6}$$

Similarly, invasion growth rate of species 2 can be written as,

$$r_2^* = 1 - \Big(\frac{\alpha_{21}(\alpha_{11} - \beta_{113}N_3^*) - \beta_{211} - \beta_{213}(\alpha_{11} - \beta_{113}N_3^*)}{(\alpha_{11} - \beta_{113}N_3^*)^2}\Big). \tag{A.7}$$

### Appendix B. Four-way higher order Lotka-Volterra model and invasion growth rates

Four way (fourth order) higher order Lotka-Volterra model can be written as :

$$\frac{dN_i}{dt} = N_i r_i(1 - \sum_j \alpha_{ij}N_j + \sum_j \sum_k \beta_{ijk}N_jN_k + \sum_j \sum_k \sum_l \gamma_{ijkl}N_jN_kN_l) \tag{B.1}$$

where, $\gamma_{ijkl}$ denotes the four-way HOIs. Specifically, $\gamma_{ijkl}$ captures the non-additive effect of species $l$ on three-way interactions of species $k$ and species $j$ on species $i$. Expanding equation 8 for species 1 yields :

$$\begin{aligned}
\frac{dN_1}{dt} = N_1r_1(1 - \alpha_{11}N_1 - \alpha_{12}N_2 + \\
\beta_{112}N_1N_2 + \beta_{113}N_1N_3 + \beta_{121}N_2N_1 + \beta_{122}N_2^2 + \beta_{123}N_2N_3 + \\
\gamma_{1121}N_1^2N_2 + \gamma_{1122}N_1N_2^2 + \gamma_{1123}N_1N_2N_3 + \gamma_{1131}N_1^2N_3 + \gamma_{1132}N_1N_3N_2 + \\
\gamma_{1133}N_1N_3^2 + \gamma_{1211}N_2N_1^2 + \gamma_{1212}N_1N_2^2 + \gamma_{1213}N_1N_2N_3 + \gamma_{1221}N_2^2N_1 + \gamma_{1222}N_2^3 + \\
\gamma_{1223}N_2^2N_3 + \gamma_{1231}N_2N_3N_1 + \gamma_{1232}N_2^2N_3 + \gamma_{1233}N_2N_3^2)
\end{aligned} \tag{B.2}$$

Typically, in a pairwise interaction Lotka-Volterra model, the number of pairwise terms in a three species model is 9. However, with the incorporation of three-way HOIs, the number of interaction terms in three way HOIs Lotka-Voterra model is 27 (considering species 3 also participates both in pairwise and HOIs). In four way HOI model, four way HOIs terms increases to 54. Henceforth, to simplify our modelling scenarios and to analyse the effect of HOIs on species coexistence, we made certain assumptions, one of which was to ensure that species 3 participates only in HOIs.

Invasion growth rate of species 1 when rare ($N_1 = 0$ in the presence of four way HOIs and in the presence of species 2 and species 3, thus can be estimated as :

$$\begin{aligned}
\bar{r}_1^* &= \frac{dN_1}{dt}\frac{1}{N_1} > 0 \\
&= 1 - \alpha_{12}N_2 + \beta_{122}N_2^2 + \beta_{123}N_2N_3 + \gamma_{1222}N_2^3 + \gamma_{1223}N_2^2N_3 + \gamma_{1232}N_2^2N_3 + \\
&\quad \gamma_{1233}N_2N_3^2 > 0,
\end{aligned} \tag{B.3}$$

where $N_2$, $N_3$ are the equilibrium abundances of species 2 and species 3. Similarly, invasion growth rate of species 2, in the presence of four way HOIs and in the presences of species 1 and species 2 can be written as :

$$\begin{aligned}
\bar{r}_2^* &= \frac{dN_2}{dt}\frac{1}{N_2} > 0 \\
&= 1 - \alpha_{21}N_1 + \beta_{211}N_1^2 + \beta_{213}N_1N_3 + \gamma_{2111}N_1^3 + \gamma_{2113}N_1^2N_3 + \gamma_{2131}N_1^2N_3 + \\
&\quad \gamma_{2133}N_1N_3^2 > 0,
\end{aligned} \tag{B.4}$$

The number of possible four way HOIs that can act on species 1 by modifying three-way and pairwise interactions are:

$$\begin{pmatrix} \gamma_{1111} & \gamma_{1112} & \gamma_{1113} \\ \gamma_{1121} & \gamma_{1122} & \gamma_{1123} \\ \gamma_{1131} & \gamma_{1132} & \gamma_{1133} \end{pmatrix}$$

In the above matrix $\gamma_{1111} =_{1112}= \gamma_{1113} = 0$ as we assumed that $\beta_{iii} = 0$, hence such higher order links are

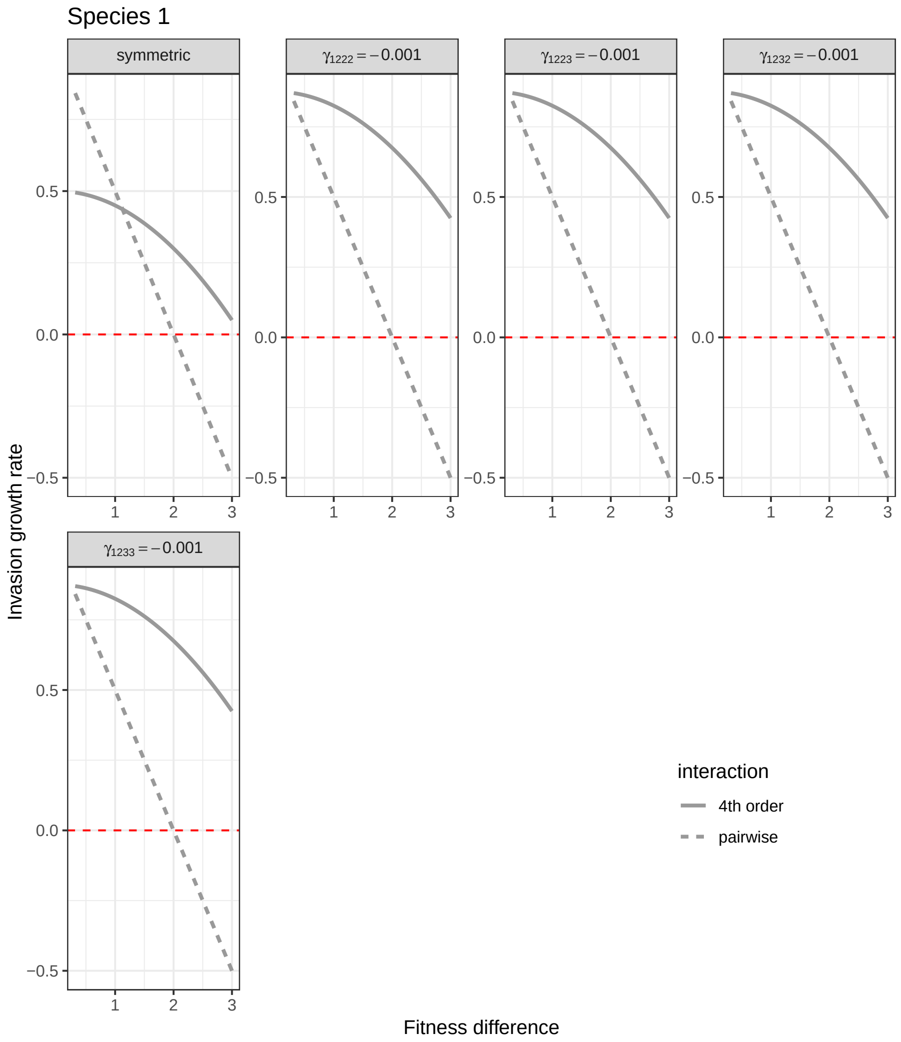


Figure B.1: Invasion growth rate of species 1, $\bar{r_1^*}$, (y-axis) for pairwise species competition (dashed grey lines) and negative four-way HOIs (grey lines) shown for a range of fitness differences (x-axis). The red-dashed line marks the y-intercept at zero. Each panel of the plot compares invasion growth rate of species 1 in pairwise competition (dashed grey lines) and in negative four-way HOIs ( solid grey continuous lines). The leftmost top panel in the first row of the figure (symmetric) denotes the case where all HOI terms are negative and have the same strength of $-0.001$. In this panel, invasion growth rate of species 1 in pure pairwise interaction with species 2 (dashed line) is plotted against invasion growth rate of species 1 in four way HOIs (grey continuous line) where all the terms of HOIs have same strength. In the other panels, invasion growth rate of species 1 in pure pairwise competition (dashed line) is plotted against invasion growth rate of species 1 in HOIs, when only one term from HOIs is perturbed, denoted by the panel labelled as $\gamma_{1222}; \gamma_{1223}; \gamma_{1232}; \gamma_{1233}$ with the rest of the terms of the four way HOI matrix being 0. For example, $\gamma_{1223} = -0.001$ would mean all the are zero except $\gamma_{1223}$ which is at -0.001.Alongside invasion growth rate of species 1 in the presence of HOIs, invasion growth rate of species 1 in pairwise competition is also plotted in each of the panels for comparison. Here $N_2 = N_3 = 5$

forbidden. The above matrix denotes all the possible four-way HOIs that could influence intraspecific pairwise interaction of species 1.

Similarly, all the four-way HOI terms that could affect interspecific pairwise interaction of species 2 on species 1 are :

$$\begin{pmatrix} \gamma_{1211} & \gamma_{1212} & \gamma_{1213} \\ \gamma_{1221} & \gamma_{1222} & \gamma_{1223} \\ \gamma_{1231} & \gamma_{1232} & \gamma_{1233} \end{pmatrix}$$

Since there are no pairwise interaction between species 1 and species 3, all possible four-way HOIs that could influence pairwise interaction of species 1 and species 3 are zero or does not exist.

Similarly, we can write the matrices of HOI terms that could influence intraspecific interaction of species 2 and interspecific interaction between 2 and 1 as:

$$\begin{pmatrix} \gamma_{2211} & \gamma_{2212} & \gamma_{2213} \\ \gamma_{2221} & \gamma_{2222} & \gamma_{2223} \\ \gamma_{2231} & \gamma_{2232} & \gamma_{2233} \end{pmatrix}$$

and,

$$\begin{pmatrix} \gamma_{2111} & \gamma_{2112} & \gamma_{2113} \\ \gamma_{2121} & \gamma_{2122} & \gamma_{2123} \\ \gamma_{2131} & \gamma_{2132} & \gamma_{2133} \end{pmatrix}$$

Here, $\gamma_{2221} = \gamma_{2222} = \gamma_{2223} = 0$

### Appendix C. Sensitivity of invasion growth rate to HOIs

Sensitivity of invasion growth rate of species 1 and species 2 to changes in strength of three way HOIs are calculated as :

$$\frac{\partial r_1^*}{\partial \beta} = \frac{\partial}{\partial \beta}\left(1 - \left(\frac{\alpha_{12}(\alpha_{22} - \beta_{223}N_3^*) - \beta_{122} - \beta_{123}(\alpha_{22} - \beta_{223}N_3^*)}{(\alpha_{22} - \beta_{223}N_3^*)^2}\right)\right) \tag{C.1}$$

and for species 2 :

$$\frac{\partial r_2^*}{\partial \beta} = \frac{\partial}{\partial \beta}\left(1 - \left(\frac{\alpha_{21}(\alpha_{11} - \beta_{113}N_3^*) - \beta_{211} - \beta_{213}(\alpha_{11} - \beta_{113}N_3^*)}{(\alpha_{11} - \beta_{113}N_3^*)^2}\right)\right) \tag{C.2}$$

Sensitivity of invasion growth rate of species 1 and species 2 to four way HOIs ( $\gamma$ terms) can similarly be calculated as above.

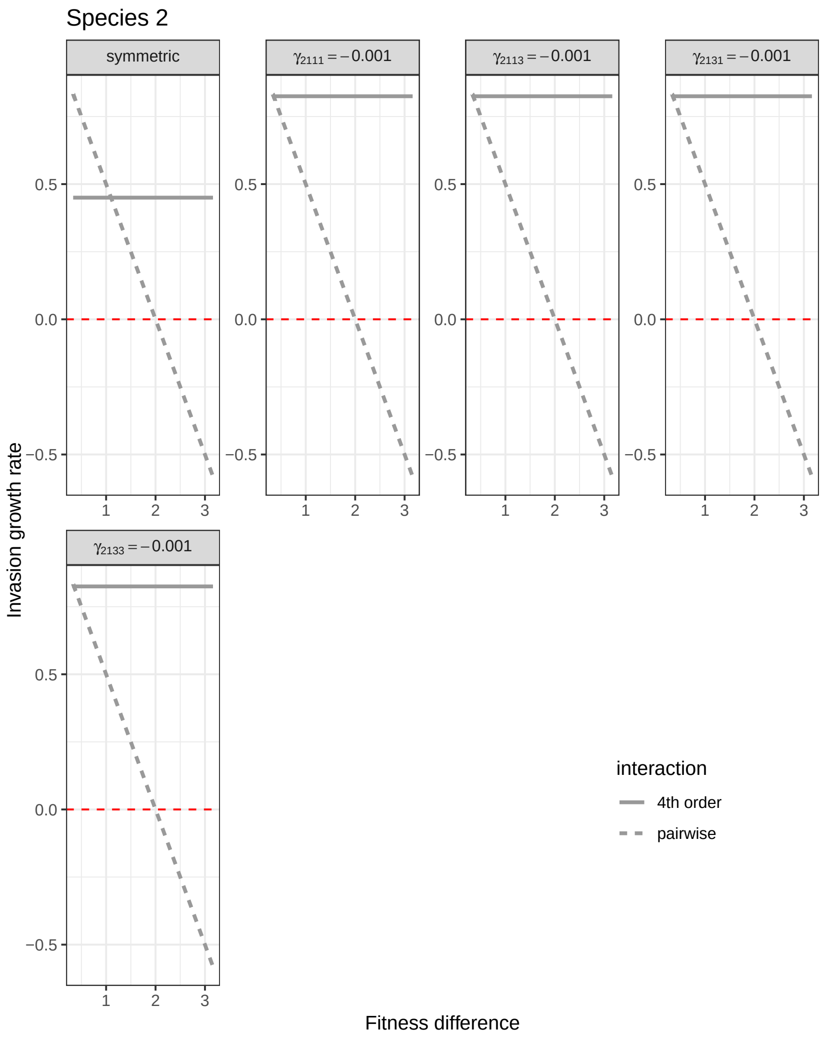


Figure B.2: Invasion growth rate of species 2, $\bar{r}_2^*$, (y-axis) for pairwise species competition (dashed grey lines) and negative four-way HOIs (grey lines) shown for a range of fitness differences (x-axis). The red-dashed line marks the y-intercept at zero. Each panel of the plot compares invasion growth rate of species 2 in pairwise competition (dashed grey lines) and in negative four-way HOIs ( solid grey continuous lines). The leftmost top panel in the first row of the figure (symmetric) denotes the case where all HOI terms are negative and have the same strength of $-0.001$. In this panel, invasion growth rate of species 2 in pure pairwise interaction with species 1 (dashed line) is plotted against invasion growth rate of species 2 in four way HOIs (grey continuous line) where all the terms of HOIs have same strength. In the other panels, invasion growth rate of species 2 in pure pairwise competition (dashed line) is plotted against invasion growth rate of species 2 in HOIs, when only one term from HOIs is perturbed, denoted by the panel labelled as $\gamma_{2111}; \gamma_{2113}; \gamma_{2131}; \gamma_{2133}$ with the rest of the terms of the HOI matrix being 0. For example, $\gamma_{2111} = -0.001$ would mean all the elements of four-way HOI matrix are zero except $\gamma_{2113}$ which is at -0.001.Alongside invasion growth rate of species 2 in the presence of HOIs, invasion growth rate of species 2 in pairwise competition is also plotted in each of the panels for comparison. Here $N_1 = N_3 = 5$.

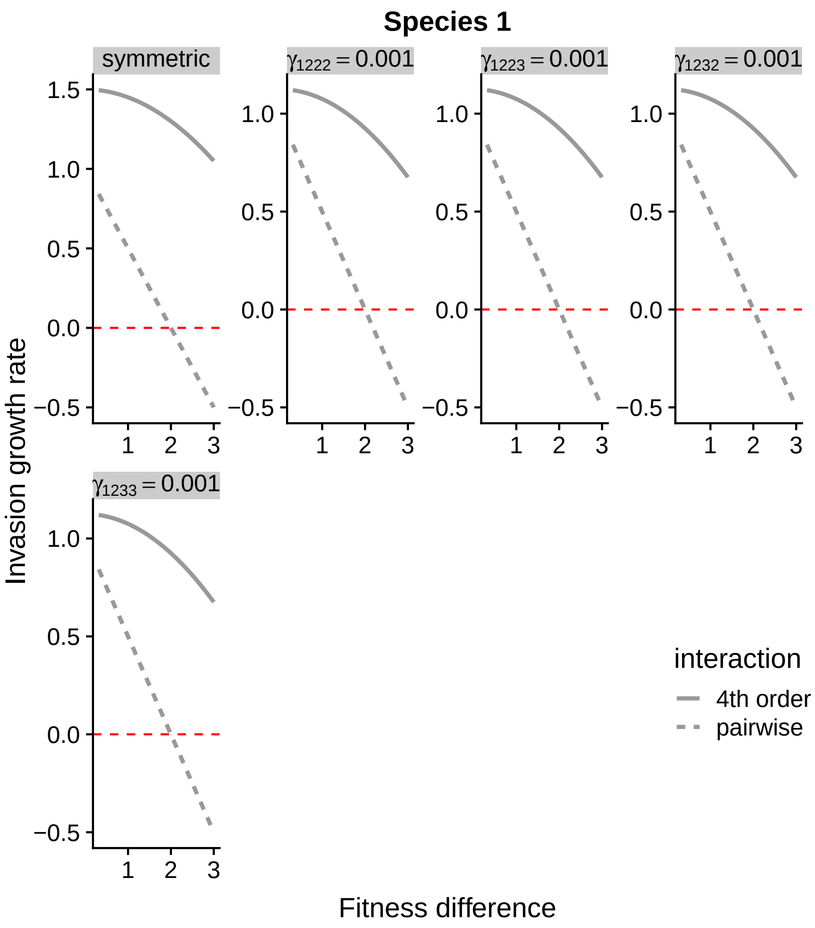


Figure B.3: Invasion growth rate of species 1, $\bar{r_1^*}$, (y-axis) for pairwise species competition (dashed grey lines) and positive four-way HOIs (grey lines) shown for a range of fitness differences (x-axis). The red-dashed line marks the y-intercept at zero. Each panel of the plot compares invasion growth rate of species 1 in pairwise competition (dashed grey lines) and in positive four-way HOIs ( solid grey continuous lines). The leftmost top panel in the first row of the figure (symmetric) denotes the case where all HOI terms are positive and have the same strength of 0.001. In this panel, invasion growth rate of species 1 in pure pairwise interaction with species 2 (dashed line) is plotted against invasion growth rate of species 1 in four way HOIs (grey continuous line) where all the terms of HOIs have same strength. In the other panels, invasion growth rate of species 1 in pure pairwise competition (dashed line) is plotted against invasion growth rate of species 1 in HOIs, when only one term from HOIs is perturbed, denoted by the panel labelled as $\gamma_{1222}; \gamma_{1223}; \gamma_{1232}; \gamma_{1233}$ with the rest of the terms of the HOI matrix being 0. For example, $\gamma_{1233} = 0.001$ would mean all the elements of four-way HOI matrix are zero except $\gamma_{1233}$ which is at 0.001.Alongside invasion growth rate of species 1 in the presence of HOIs (solid grey lines), invasion growth rate of species 2 in pairwise competition (dashed line) is also plotted in each of the panels for comparison. Here $N_2 = N_3 = 5$

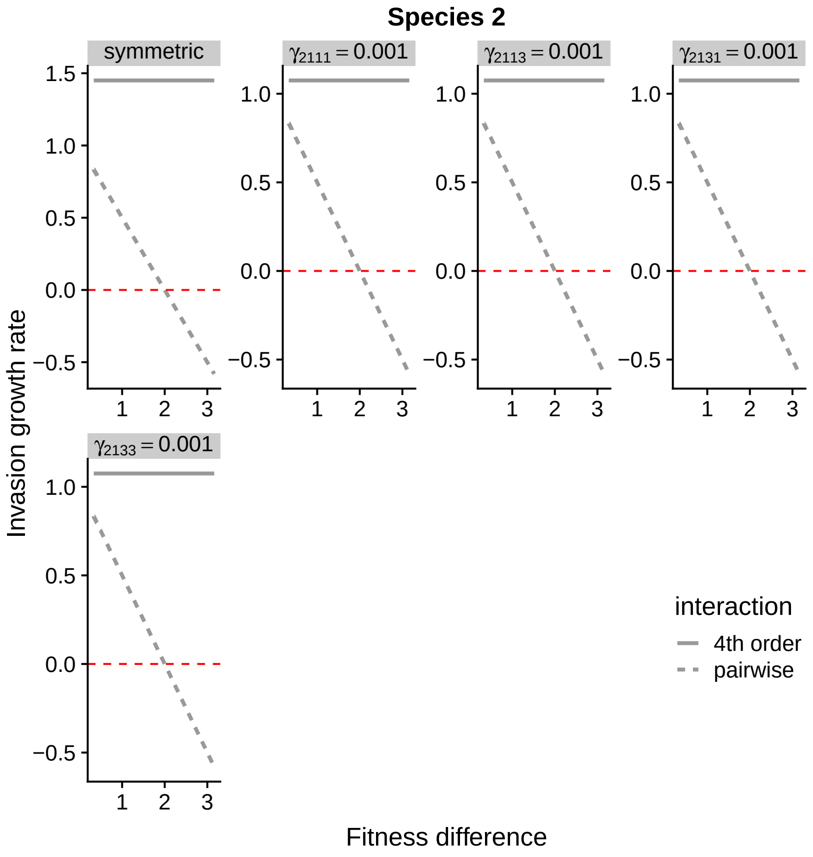


Figure B.4: Invasion growth rate of species 2, $\bar{r}_2^*$, (y-axis) for pairwise species competition (dashed grey lines) and positive four-way HOIs (grey lines) shown for a range of fitness differences (x-axis). The red-dashed line marks the y-intercept at zero. Each panel of the plot compares invasion growth rate of species 2 in pairwise competition (dashed grey lines) and in positive four-way HOIs ( solid grey continuous lines). The leftmost top panel in the first row of the figure (symmetric) denotes the case where all HOI terms are positive and have the same strength of 0.001. In this panel, invasion growth rate of species 2 in pure pairwise interaction with species 1 (dashed line) is plotted against invasion growth rate of species 2 in four way HOIs (grey continuous line) where all the terms of HOIs have same strength. In the other panels, invasion growth rate of species 2 in pure pairwise competition (dashed line) is plotted against invasion growth rate of species 2 in HOIs, when only one term from HOIs is perturbed, denoted by the panel labelled as $\gamma_{2111}; \gamma_{2113}; \gamma_{2131}; \gamma_{2133}$ with the rest of the terms of the HOI matrix being 0. For example, $\gamma_{2111} = -0.001$ would mean all the elements of four-way HOI matrix are zero except $\gamma_{2113}$ which is at -0.001.Alongside invasion growth rate of species 2 in the presence of HOIs, invasion growth rate of species 2 in pairwise competition is also plotted in each of the panels for comparison. Here $N_1 = N_3 = 5$

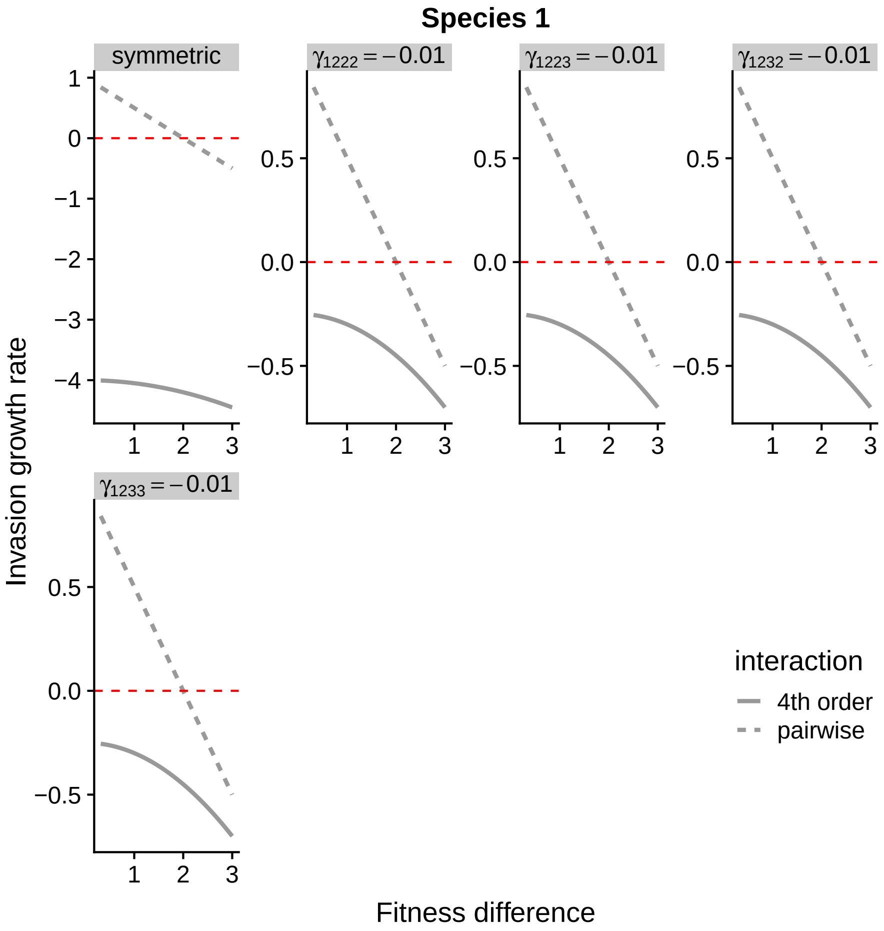


Figure B.5: Effect of stronger four-way HOIs on species coexistence. Invasion growth rate of species 1 $\bar{r_1^*}$ (y-axis) for pairwise species competition (dashed grey lines) and negative four-way HOIs (grey lines) shown for a range of fitness differences (x-axis). The red-dashed line marks the y-intercept at zero. Each panel of the plot compares invasion growth rate of species 1 in pairwise competition (dashed grey lines) and in negative four-way HOIs ( solid grey continuous lines). The leftmost top panel in the first row of the figure (symmetric) denotes the case where all HOI terms are negative and have the same strength of $-0.01$. In this panel, invasion growth rate of species 1 in pure pairwise interaction with species 2 (dashed line) is plotted against invasion growth rate of species 1 in four way HOIs (grey continuous line) where all the terms of HOIs have same strength. In the other panels, invasion growth rate of species 1 in pure pairwise competition (dashed line) is plotted against invasion growth rate of species 1 in HOIs, when only one term from HOIs is perturbed, denoted by the panel labelled as $\gamma_{1222}; \gamma_{1223}; \gamma_{1232}; \gamma_{1233}$ with the rest of the terms of the HOI matrix being 0. For example, $\gamma_{1223} = -0.01$ would mean all the elements of four-way HOI matrix are zero except $\gamma_{1223}$ which is at -0.01. Alongside invasion growth rate of species 1 in the presence of HOIs, invasion growth rate of species 1 in pairwise competition is also plotted in each of the panels for comparison. Note that as strength of four-way HOIs increased from -0.001 to -0.01, coexistence was impossible regardless of negligible fitness differences when HOIs are prevalent. Here $N_2 = N_3 = 5$

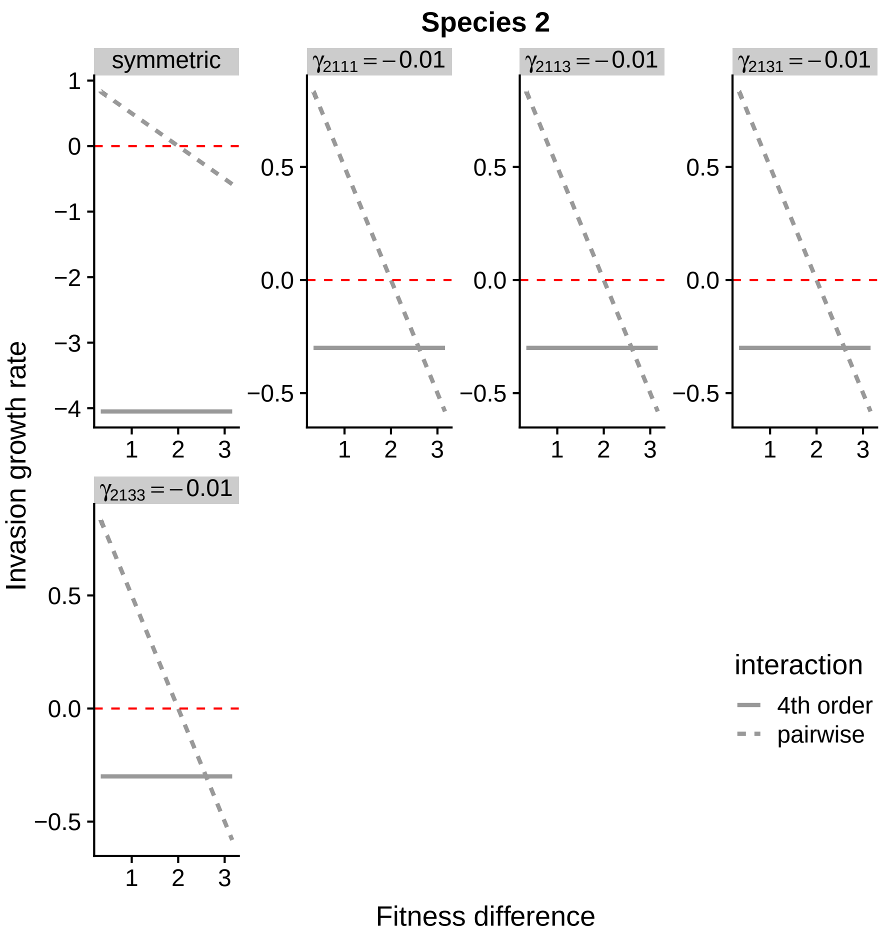


Figure B.6: Invasion growth rate of species 2, $\bar{r_2^*}$, (y-axis) for pairwise species competition (dashed grey lines) and negative four-way HOIs (grey lines) shown for a range of fitness differences (x-axis). The red-dashed line marks the y-intercept at zero. Each panel of the plot compares invasion growth rate of species 2 in pairwise competition (dashed grey lines) and in negative four-way HOIs (solid grey lines). The first leftmost panel on in the left top row of the figure (symmetric) denotes the case where all HOI terms are negative and have the same strength of $-0.01$. In the other panels, only one term from is perturbed, denoted by the panel labelled as $\gamma_{2111}; \gamma_{2113}; \gamma_{2131}; \gamma_{2133}$ with the rest of the terms of the HOI matrix being 0. For example, $\gamma_{2111} = -0.01$ would mean all the elements of four-way HOI matrix are zero except $\gamma_{2111}$ which is at -0.01. Alongside invasion growth rate of species 2 in the presence of HOIs, invasion growth rate of species 2 in pairwise competition is also plotted in each of the panels for comparison.Note that as strength of four-way HOIs increased from -0.001 to -0.01, coexistence was impossible regardless of negligible fitness differences when HOIs are prevalent. Here $N_1 = N_3 = 5$

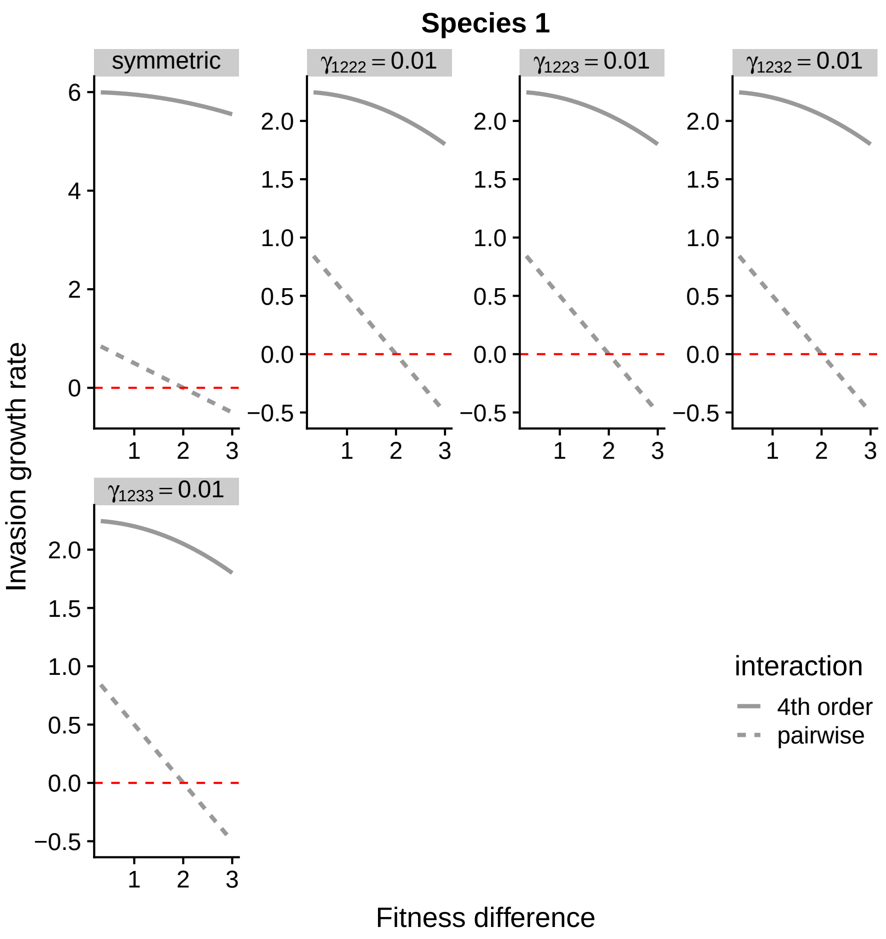


Figure B.7: Effect of stronger four-way HOIs on species coexistence. Invasion growth rate of species 1, $\bar{r_1^*}$ , (y-axis) for pairwise species competition (dashed grey lines) and positive four-way HOIs (grey lines) shown for a range of fitness differences (x-axis). The red-dashed line marks the y-intercept at zero. Each panel of the plot compares invasion growth rate of species 1 in pairwise competition (dashed grey lines) and in positive four-way HOIs ( solid grey continuous lines). The leftmost top panel in the first row of the figure (symmetric) denotes the case where all HOI terms are positive and have the same strength of 0.01. In this panel, invasion growth rate of species 1 in pure pairwise interaction with species 2 (dashed line) is plotted against invasion growth rate of species 1 in four way HOIs (grey continuous line) where all the terms of HOIs have same strength. In the other panels, invasion growth rate of species 1 in pure pairwise competition (dashed line) is plotted against invasion growth rate of species 1 in HOIs, when only one term from HOIs is perturbed, denoted by the panel labelled as $\gamma_{1222}; \gamma_{1223}; \gamma_{1232}; \gamma_{1233}$ with the rest of the terms of the HOI matrix being 0. For example, $\gamma_{1223} = 0.01$ would mean all the elements of four-way HOI matrix are zero except $\gamma_{1223}$ which is at 0.01. Alongside invasion growth rate of species 1 in the presence of HOIs, invasion growth rate of species 1 in pairwise competition is also plotted in each of the panels for comparison. Note that as strength of positive four-way HOIs increased from 0.001 to 0.01, coexistence was always regardless of high fitness differences when HOIs are prevalent. Positive four way HOIs leads to lessening of pairwise and three-way interactions. Here $N_2 = N_3 = 5$

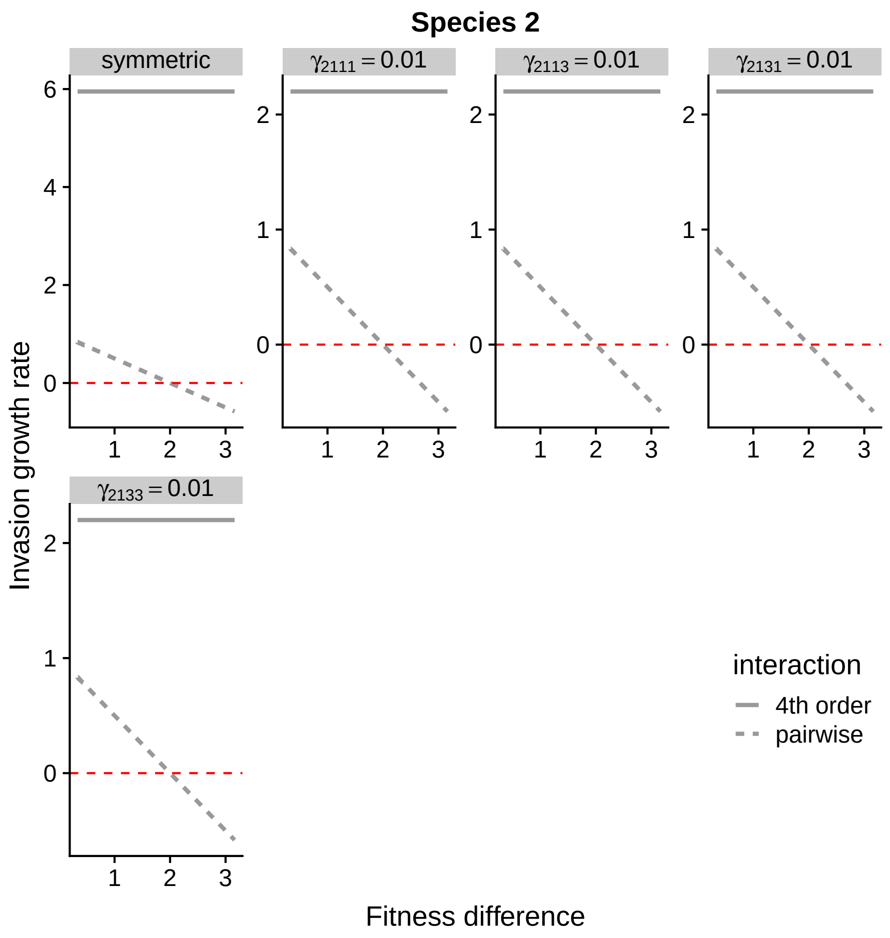


Figure B.8: Invasion growth rate of species 2, $\bar{r_2^*}$ , (y-axis) for pairwise species competition (dashed grey lines) and positive four-way HOIs (grey lines) shown for a range of fitness differences (x-axis). The red-dashed line marks the y-intercept at zero. Each panel of the plot compares invasion growth rate of species 2 in pairwise competition (dashed grey lines) and in positive four-way HOIs (solid grey lines). The first leftmost panel on in the left top row of the figure (symmetric) denotes the case where all HOI terms are positive and have the same strength of 0.01. In the other panels, only one term from is perturbed, denoted by the panel labelled as $\gamma_{2111}; \gamma_{2113}; \gamma_{2131}; \gamma_{2133}$ with the rest of the terms of the HOI matrix being 0. For example, $\gamma_{2111} = 0.01$ would mean all the elements of HOI matrix are zero except $\gamma_{2111}$ which is at 0.01. Alongside invasion growth rate of species 2 in the presence of HOIs, invasion growth rate of species 2 in pairwise competition is also plotted in each of the panels for comparison.Note that as strength of four-way HOIs increased from 0.001 to 0.01, coexistence was always possible regardless of high fitness differences when HOIs are prevalent. Here $N_1 = N_3 = 5$

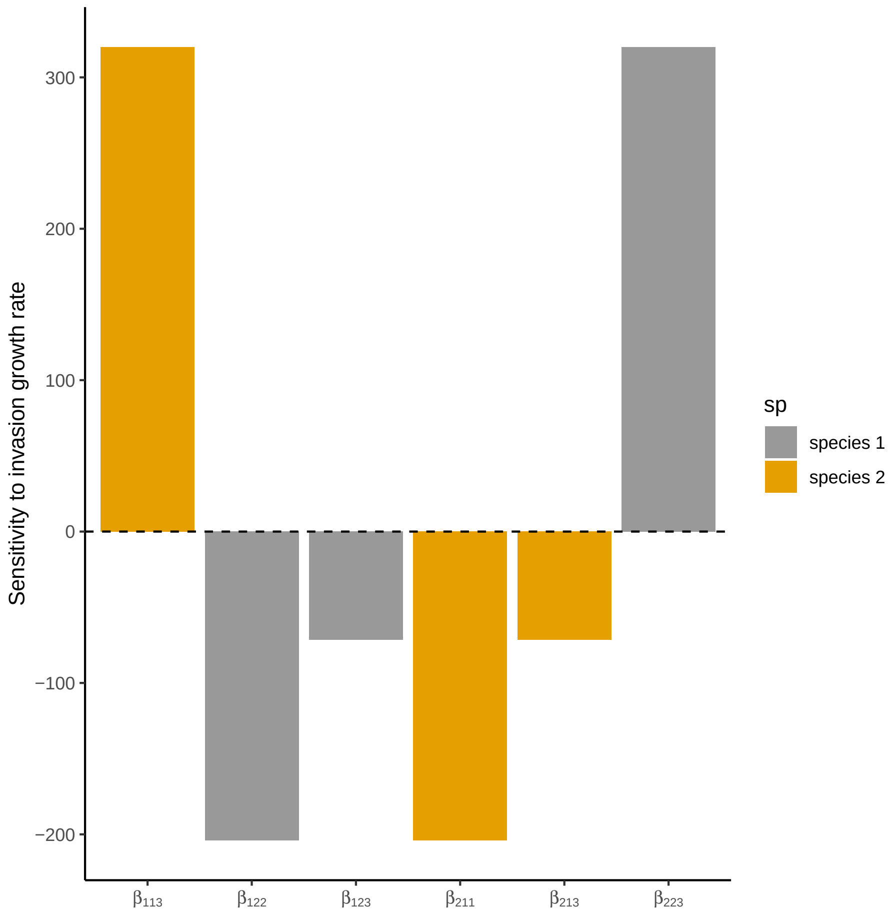


Figure C.9: Sensitivity of invasion growth rate of species 1 and species 2 to three-way HOIs. Note that invasion growth rate of species 1 and species 2 will increase and hence stabilize coexistence only when $\beta_{113}$ or $\beta_{223}$ increases more than other HOI terms. In order words, if intraspecific pairwise competition is strengthened more by three-way interactions, species coexistence is stabilized.

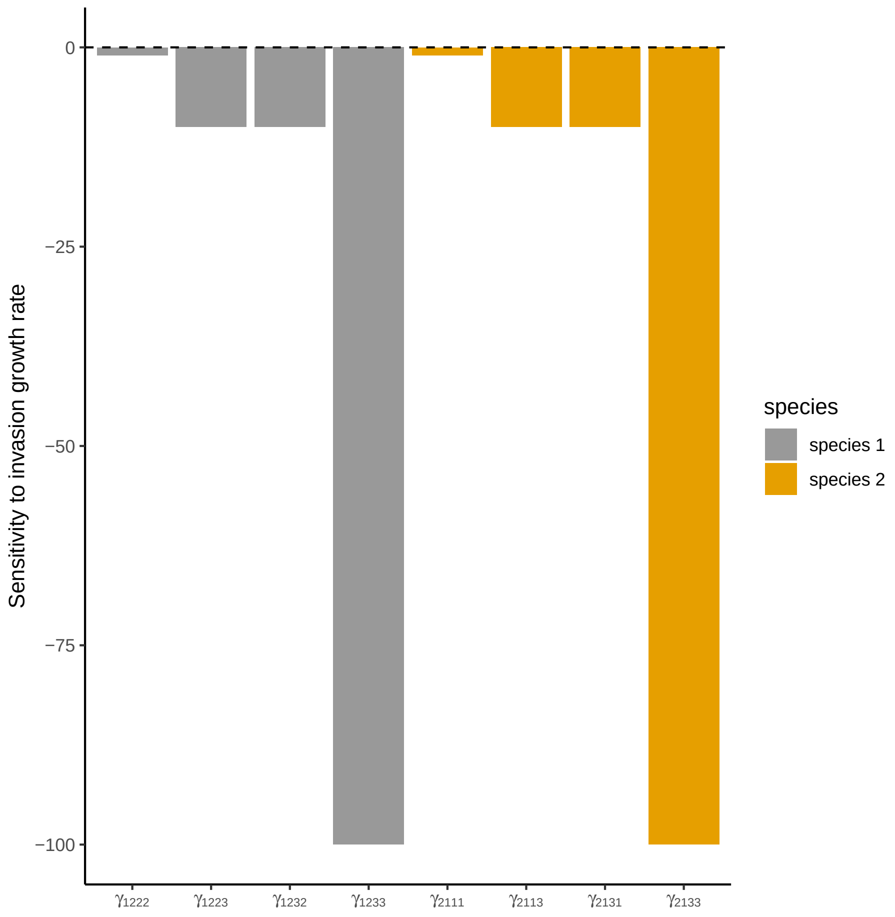


Figure C.10: Sensitivity of invasion growth rate of species 1 and species 2 to three-way HOIs. Note that invasion growth rate of species 1 and species 2 will always be less than zero and hence destabilize coexistence any of the fourth-order interspecific HOIs increases more.

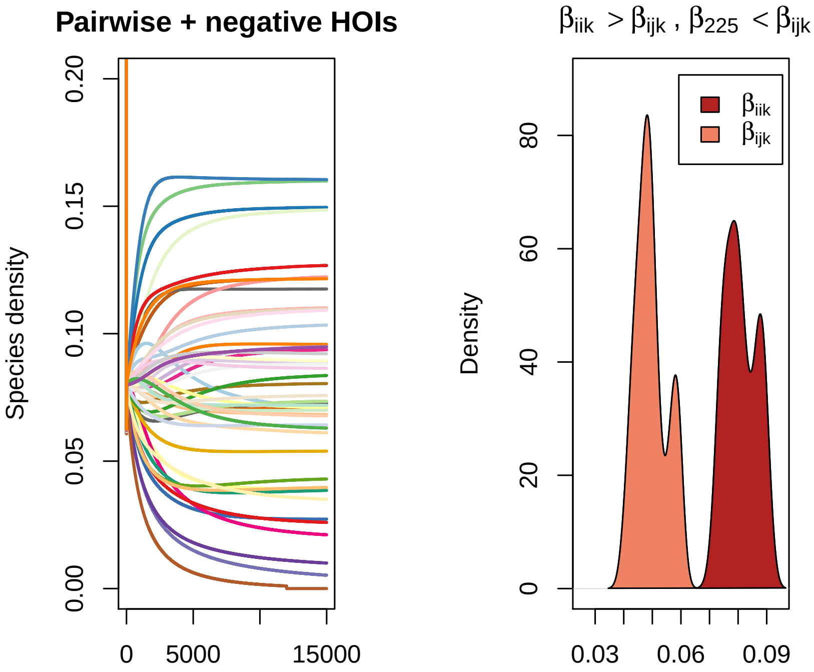


Figure C.11: In the above figure, Weyl's inequality not satisfied which means pairwise species coexistence is not possible. However, when intraspecific HOIs $\beta_{iik} > \beta_{ijk}$ species coexistence is possible. But if one HOI interaction does not fullfill the criteria for multispecies coexistence rule, that intraspecific HOIs should be greater than interspecific HOIs, $\beta_{225} < \beta_{ijk}$, coexistence of all 50 species was not possible and species 2 went extinct.

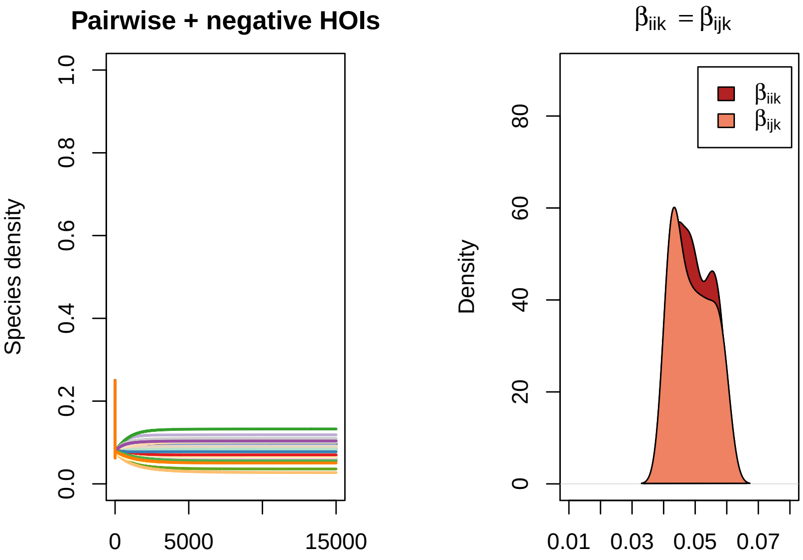


Figure C.12: Weyl's inequality IS satisfied which means pairwise species coexistence is always possible. However, when intraspecific HOIs $\beta_{iik}$ $\beta_{ijk}$, i.e. the distribution of intraspecific HOIs overlaps with distribution of interspecific HOIs, species coexistence is still possible.

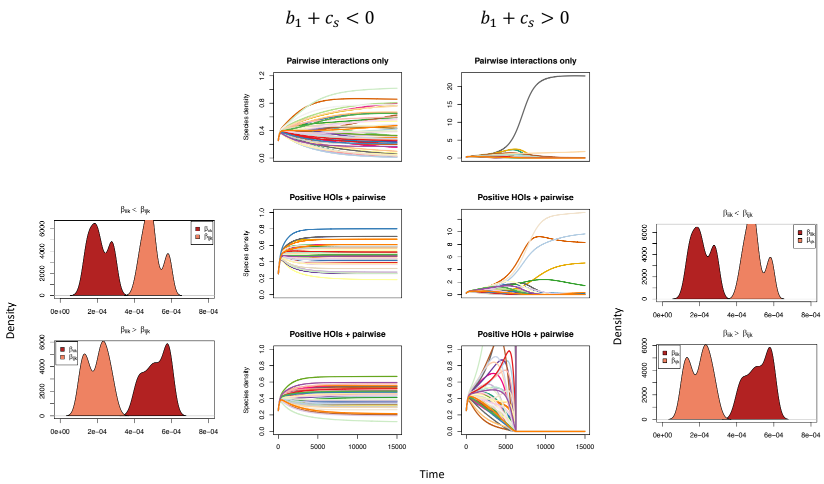


Figure C.13: Positive HOIs on multispecies coexistence either satisfying Weyl's inequality or not. When Weyl's inequality is satisfied $b_1 + c_s < 0$, and there is strong self-regulation, positive HOIs can stabilize species coexistence. Pairwise coexistence is however also possible when Weyl's inequality is satisfied. When Weyl's inequality is not satisfied $b_1 + c_s > 0$, species coexistence is disrupted both by pairwise interactions and positive HOIs.

### Appendix D. References


[1] Baruah, G. and R. John (2018, 12). Intraspecific variation promotes trait clustering and species coexistence through higher order interactions. *bioRxiv*, 494757.

[2] Grilli, J., G. Barabás, M. J. Michalska-Smith, and S. Allesina (2017, 7). Higher-order interactions stabilize dynamics in competitive network models. *Nature 548*(7666), 210.

[3] Hinsinger, P., A. G. Bengough, D. Vetterlein, and I. M. Young (2009, 8). Rhizosphere: biophysics, biogeochemistry and ecological relevance. *Plant and Soil 321*(1-2), 117–152.

[4] Mayfield, M. M. and D. B. Stouffer (2017, 2). Higher-order interactions capture unexplained complexity in diverse communities. *Nature Ecology & Evolution 1*(3), 0062.

[5] Terry, J. C. D., R. J. Morris, and M. B. Bonsall (2017, 10). Trophic interaction modifications: an empirical and theoretical framework. *Ecology Letters 20*(10), 1219–1230.
